# Genotype-specific Features Reduce the Susceptibility of South American Yellow Fever Virus Strains to Vaccine-Induced Antibodies

**DOI:** 10.1101/2021.08.22.457235

**Authors:** Denise Haslwanter, Gorka Lasso, Anna Z. Wec, Nathália Dias Furtado, Lidiane Menezes Souza Raphael, Yan Sun, Stephanie Stransky, Núria Pedreño-Lopez, Alexandra Tse, Carolina Argondizo Correia, Zachary A. Bornholdt, Mrunal Sakharkar, Vivian I. Avelino-Silva, Crystal L. Moyer, David I. Watkins, Esper G. Kallas, Simone Sidoli, Laura M. Walker, Myrna C. Bonaldo, Kartik Chandran

## Abstract

The resurgence of yellow fever in South America has prompted mitigation through vaccination against the etiologic agent, yellow fever virus (YFV). Current vaccines are based on a virulent African isolate, and their capacity to induce neutralizing antibodies against the vaccine strain is widely used as a surrogate for protection. However, the sensitivity of genetically distinct South American strains to vaccine-induced antibodies is unknown. Here, we show that antiviral potency of the polyclonal antibody response in both U.S. and Brazilian vaccinees is attenuated against an emergent Brazilian strain. This reduction was attributable to genetic changes at two sites in the central domain II of the glycoprotein E, including the acquisition of an *N*–linked glycosylation site, which are unique to and shared among most South American YFV strains. Our findings call for a reevaluation of current approaches to YFV immunological surveillance in South America and suggest approaches for designing updated vaccines.

## Introduction

Yellow fever virus (YFV), a prototypic flavivirus and the etiologic agent of yellow fever, is endemic to tropical and subtropical regions of Africa, Central and South America. Although YFV largely circulates in a sylvatic cycle between mosquitoes and nonhuman primates, mosquito-borne viral transmission from nonhuman primates to humans can occur due to encroachment by the latter into forested areas. Further, high population densities in urban areas can trigger cycles of human-to-human transmission that are maintained by anthropophilic mosquitoes. YFV has periodically resurged in endemic areas in South America since the 1960s, but especially intense reemergence events have been documented in both endemic and non-endemic areas in the past decade (Douam and Ploss, 2018). The 2016–19 YFV epidemic centered in southeastern Brazil was the largest in over 70 years, with >2,000 cases and >750 deaths (Hill et al., 2020; de Oliveira Figueiredo et al., 2020).

The live-attenuated YF-17D virus was derived from the virulent African strain Asibi and was further passaged to yield closely related variants (e.g., YF-17D-204 and YF-17DD; hereafter, YF-17D) that form the basis of YFV vaccines currently in use. These are among the most effective vaccines ever created and remain the linchpin of global efforts to control yellow fever (Barrett and Teuwen, 2009). Single doses of YF-17D are expected to confer protection for at least 2–3 decades post-immunization (Barrett and Teuwen, 2009; Monath, 2012). The drivers of YFV reemergence in South America and elsewhere in the face of YF-17D vaccination are likely complex and related to multiple factors, including ecological disruptions attendant to human activity and climate change, inadequate mosquito control, and suboptimal vaccination rates exacerbated by ongoing vaccine shortages (WHO, 2018). An understanding of the factors that influence YFV reemergence and vaccine efficacy is crucial for the prevention and management of future outbreaks.

Current evidence indicates that the 2017-19 YFV epidemic in Brazil was associated with an emerging strain, YFV 2017-19, which bears multiple nonsynonymous changes at conserved sequences in several nonstructural proteins (Bonaldo et al., 2017; Giovanetti et al., 2019; Gómez et al., 2018; Silva et al., 2020). These observations prompted hypotheses that one or more viral genome sequence polymorphisms could influence YFV emergence, transmission, and/or virulence (Bonaldo et al., 2017). However, previous studies have not examined the alternative or additional possibility that antiviral activity of the vaccine-induced antibody response could be affected by genetic changes in circulating YFV strains.

The induction of neutralizing, or viral entry-blocking, antibodies targeting the envelope protein, E, is considered the most important surrogate marker for vaccine-induced protection and has been described as the major mechanism that confers long-lasting protection against flavivirus disease (Mason et al., 1973; Plotkin, 2010). Here, we show that the neutralization potency of the YFV vaccine-induced human polyclonal antibody and monoclonal antibody responses against the emergent, and now endemic, YFV 2017-19 strain is substantially lower than that predicted from classical potency assays employing the vaccine strains or a wild-type African YFV strain. This reduction was largely attributable to genetic changes at two sites in the central domain II of the YFV 2017-19 E protein. Our phylogenetic analysis of available YFV genome sequences revealed that these changes are completely absent in all African and Asian (ex-African) sequences but shared among essentially all South American sequences dating back to 1977. Finally, analysis of a large panel of vaccinee-derived E-directed monoclonal antibodies (mAbs) pointed to the existence of currently undefined antibody specificities that contribute to viral neutralization by vaccinee sera in a manner that is sensitive to sequence polymorphisms in the domain I–domain II hinge. Our findings have potential implications for understanding the molecular basis of YFV’s adaptation to sylvatic cycles following its introduction into the Americas, managing outbreaks through YF-17D vaccination in South America, and designing updated YFV vaccines.

## Results

### The Brazilian isolate YFV-ES-504 is poorly neutralized by YFV-17D vaccinee sera

We first examined the breadth of the anti-YFV neutralizing antibody response over time in three previously described U.S. YF-17D vaccinees (donors 1–3) with no serological evidence of prior flavivirus exposure and well-characterized mAbs responses (Wec et al., 2020) **(Table S1)**. Neutralizing activity in sera from the U.S. donors was measured at days 0, 10, 14, 16, 28, 90, 180, 270 and 360 post vaccination **(Figure 1A–B)** with a widely employed West Nile virus-based reporter viral particle system (RVP) (Pierson et al., 2006). RVPs faithfully recapitulate the behavior of authentic flaviviruses in studies of virus structure, function, and antibody recognition and neutralization (Bohning et al., 2021; Bradt et al., 2019; Dowd et al., 2011; Goo et al., 2016, 2017; Mattia et al., 2011) **(Figure S1)**. Serum neutralizing titers against RVPs bearing the structural proteins C-prM-E from the YF-17D vaccine strain (RVP_17D_) and the virulent African strains Asibi (RVP_Asibi_) and China (ex-Angola) (RVP_China_) **(Figure 1A)** rose sharply, peaked at days 14–16, and remained stable thereafter, as reported previously (Lindsey et al., 2018; Wec et al., 2020; Wieten et al., 2016). By contrast, we observed little or no increase in neutralizing titers against RVP_ES-504_, bearing E from the prototypic YFV 2017-19 isolate YFV-ES-504 (ES-504/BRA/2017) (Bonaldo et al., 2017) over days 10-16 in all three donors, and the titers remained at ∼10-fold lower levels relative to RVP_17D_ at all timepoints tested **(Figure 1A–B)**.

**Figure 1.**
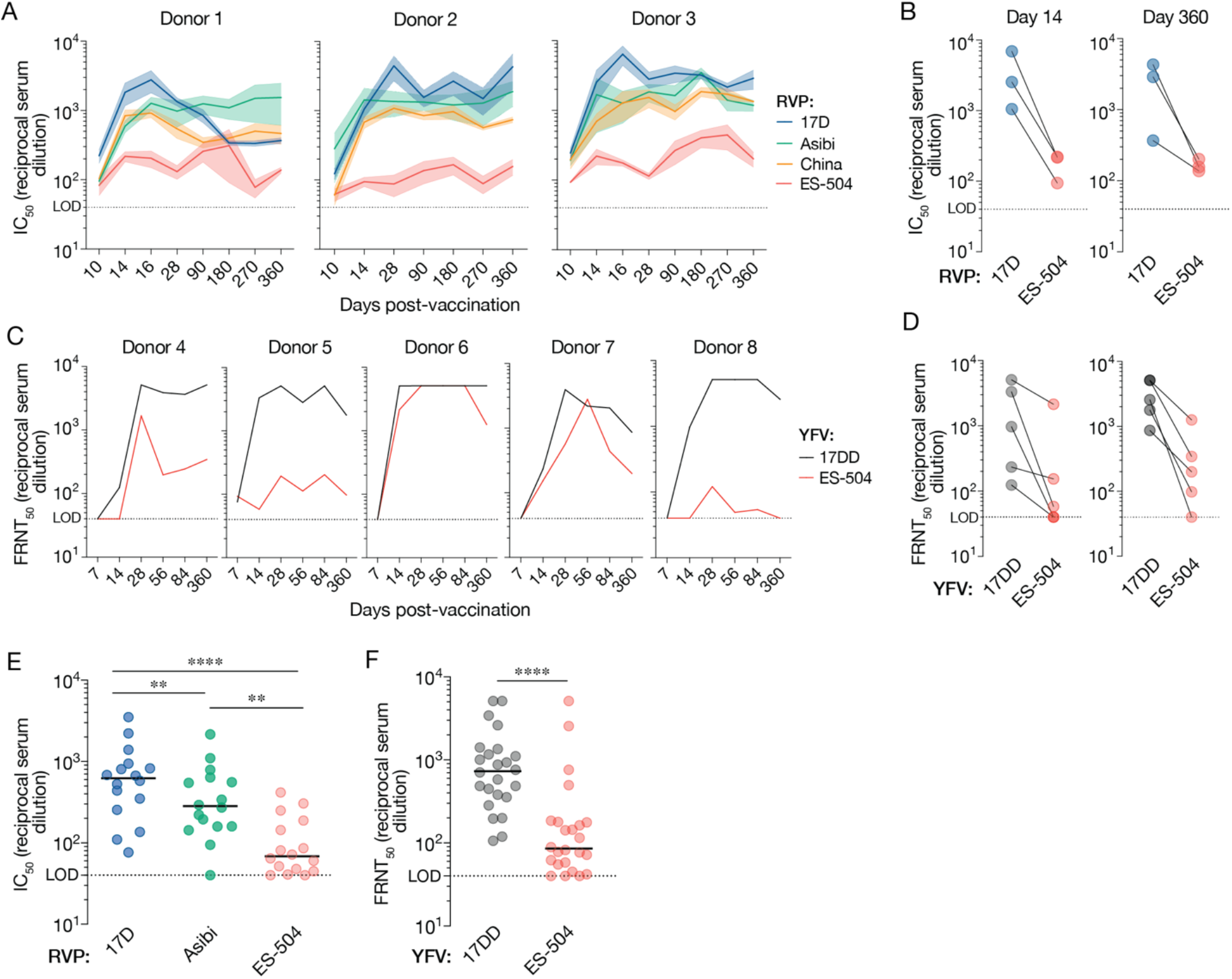
Antiviral potency and breadth of the neutralizing antibody response elicited by YF-17D vaccination. (**A**) Serum neutralizing titers (half-maximal inhibitory concentration values from dose-response curves [IC_50_]) for three YF-17D–vaccinated U.S. donors over time against the indicated RVPs. Means±SEM, n=9–15 from 3-5 independent experiments. (**B**) Serum neutralizing titers for the three donors in panel A at days 14 and 360 post-vaccination. Means, n=9–15 from 3-5 independent experiments. (**C**) Serum neutralizing titers (half-maximal inhibitory concentration values in a focus reduction neutralization test [FRNT_50_]) for five YF-17DD– vaccinated Brazilian donors over time against the indicated authentic viruses. Means, n=3. (**D**) Serum neutralizing titers (FRNT_50_) for the five donors in panel C at days 14 and 360 post-vaccination. Means, n=3. (**E**) Serum neutralizing titers for a U.S. vaccinee cohort (n=16 donors) against the indicated RVPs. n=9–12 from 3–4 independent experiments. (**F**) Serum neutralizing titers (FRNT_50_) for a Brazilian vaccinee cohort (n=24 donors against the indicated authentic viruses. n=3. In panels E–F, lines indicate group medians. Groups in E were compared by two-way ANOVA followed by Tukey’s correction for multiple comparisons. Groups in F were compared by the Wilcoxon matched-pairs signed-rank test. **, P<0.002. ****, P<0.0001. LOD, limit of detection.

We next assessed the neutralizing antibody response to YF-17D vaccination in five Brazilian donors (donors 4–8) with unknown flavivirus exposure history at days 0, 7, 14, 28, 56, 84 and 360 post vaccination **(Figure 1C and Table S1)**. Comparison of serum neutralizing titers against authentic YFV-17D and YFV-ES-504 yielded results resembling those observed in the U.S. vaccinees: blunted increases in YFV-ES-504 neutralizing activity at early times post-vaccination and substantially reduced neutralization potency relative to the vaccine strain, especially at later times in three of five donors **(Figure 1C-D)**. Donor sera from larger cohorts of U.S. and Brazilian vaccinees **(Figures 1E–F and Table S1)** corroborated this trend and revealed statistically significant reductions in YFV-ES-504 neutralization relative to YF-17D. Together, these findings indicate that vaccination with YF-17D elicits an antibody response that is substantially less effective at *in vitro* neutralization of the Brazilian YFV-ES-504 isolate than of the vaccine strains.

### Reduced YFV-ES-504 recognition and neutralization by a panel of mAbs isolated from YF-17D vaccinees

We previously isolated and characterized a large collection of YFV E-specific mAbs from U.S. YF-17D vaccinees (donors 1-3) (Wec et al., 2020). To further investigate the antiviral breadth of the vaccine-elicited antibody response, we screened 105 mAbs from this collection for their capacity to recognize the ES-504 E protein and neutralize viral particles bearing it. We measured large reductions in nAb binding to a recombinant YFV-ES-504 E protein relative to YFV-Asibi E by biolayer interferometry (BLI) **(Figure 2A)**. Similar trends in loss of YFV-ES-504 neutralization potency were observed in neutralization assays with RVPs **(Figure 2B)** and the authentic viruses **(Figure 2C)**.

**Figure 2.**
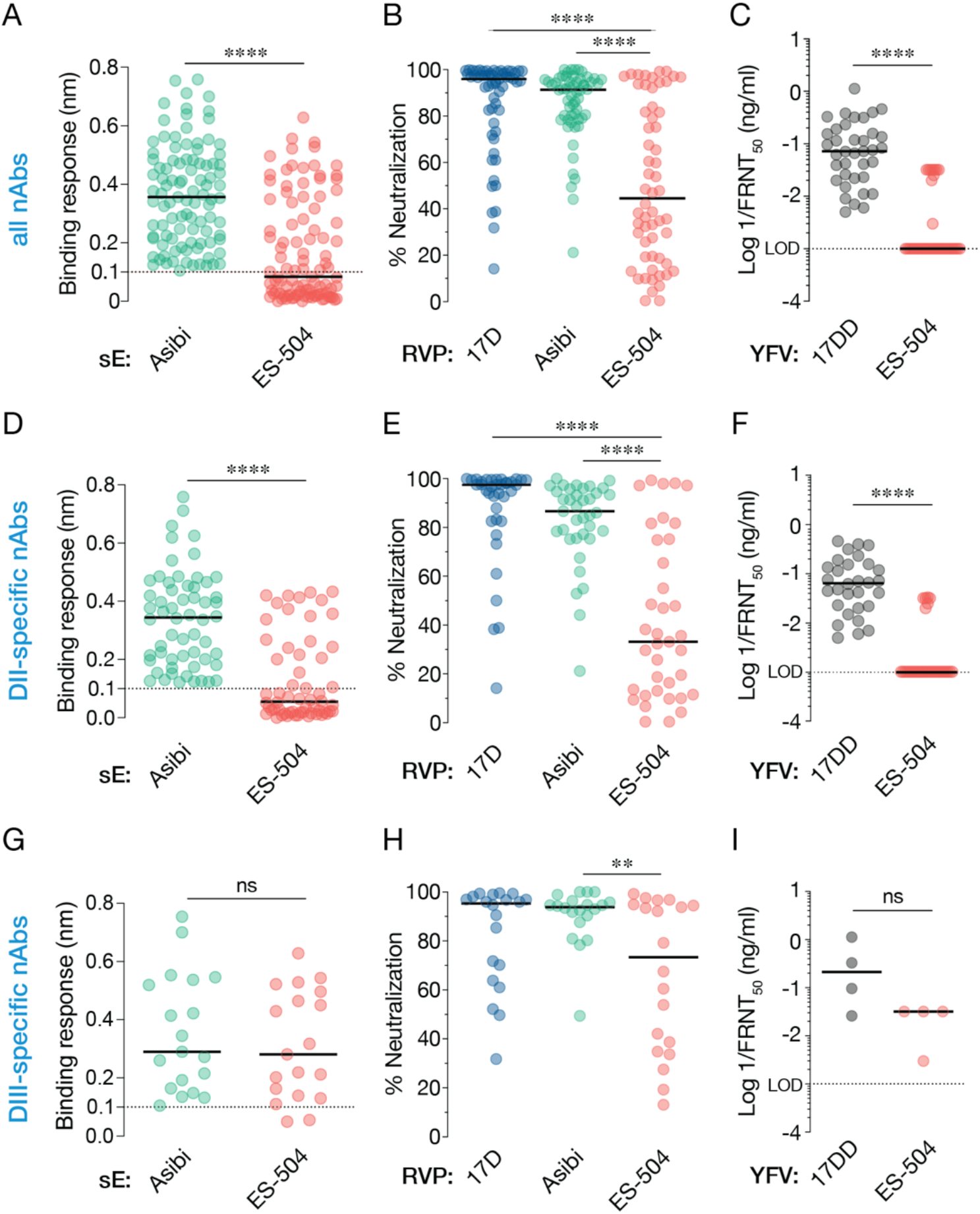
Binding and neutralization breadth of a panel of nAbs isolated from YF-17D vaccinees. (**A**) 99 nAbs were tested for their binding response to recombinant, soluble E (sE) proteins from the indicated viruses by biolayer interferometry. (**B**) Neutralizing activities of selected nAbs (10 nM) against the indicated RVPs. Means, n=6–9 from three independent experiments, (**C**) Neutralizing activities of selected mAbs against the indicated authentic viruses. (**D**) A subset of nAbs comprising DII binders were evaluated for sE binding as in panel A. (**E–F**) Neutralizing activities of DII-specific nAbs against the indicated RVPs (**E**) and authentic viruses **(F)**, as in panels B–C. (**G–I**) Binding (**G**) and neutralization activities (**H–I**) of DIII-specific nAbs, as in panels B–C. In all panels, lines indicate group medians. Groups in A, C, D, F, G, and I were compared by the Wilcoxon matched-pairs signed-rank test. Groups in B, E, H were compared by two-way ANOVA followed by Tukey’s correction for multiple comparisons. **, P<0.002. ****, P<0.0001. ns, not significant. Only the significant comparisons are shown in panels B, E, H. LOD, limit of detection.

The major surface protein E of YFV resembles those of other flavivirus envelope glycoproteins in comprising three domains, domain I, II and III (hereafter, DI, DII and DIII, respectively). In the mature virion, 90 head-to-tail dimers of E are organized into a T=3 icosahedral capsid that packages the viral genome and mediates its cytoplasmic delivery (Rey et al., 2018). A majority of the nAbs in our panel recognize non-fusion–loop antigenic sites in domain II (DII) of the YFV-17D E protein, whereas a significant minority recognize domain III (DIII) (Wec et al., 2020). To begin to define the molecular basis of reduced YFV-ES-504 binding and neutralization by YF-17D vaccinee antibodies, we curated subsets of potent DII- and DIII-specific nAbs and screened them for binding and neutralization, as above **(Figure 2D–I and Supplementary Excel File 1)**. We found that the YFV-ES-504–dependent reductions in nAb:E binding could be largely recapitulated by the DII nAbs alone **(Figure 2D and G)**. Concordantly, the DII nAbs suffered substantial reductions in neutralizing potency against viral particles bearing YFV-ES-504 E **(Figure 2E–F)**, whereas the DIII nAbs were only moderately affected **(Figure 2H–I)**. These findings raised the possibility that antibodies in human vaccinee sera specific for DII, and to a lesser degree, DIII, could significantly influence the capacity of these sera to neutralize YFV-ES-504 *in vitro* **(Figure 1)**.

### Sequence polymorphisms in domain II of the glycoprotein E largely account for the reduced neutralization sensitivity of YFV-ES-504

To uncover the genetic basis of this viral strain-dependent difference in neutralization sensitivity, we next generated and tested RVPs bearing a panel of E chimeras in which DI, DII, or DIII were exchanged between YFV-17D and YFV-ES-504 **(Figure 3A)**. RVP_17D_ bearing YFV-ES-504 DII were much less sensitive to neutralization by the DII-specific nAbs, but remained susceptible to the DIII-specific nAbs, as expected **(Figure S2)**. Importantly, these DII-chimeric RVPs also showed reduced sensitivity to neutralization by the U.S. vaccinee sera relative to wildtype (WT) RVP_17D_ at all three timepoints tested **(Figure 3B)**. Conversely, the reciprocal RVP_ES-504_ chimera bearing YFV-17D DII showed enhanced neutralization sensitivity relative to its parent **(Figure 3C)**. Analysis of a larger cohort of U.S. YFV-17D vaccinees (n=16; see **Figure 1E**) corroborated these trends: we measured a ∼10-fold reduction in median neutralization titer with RVP_17D_–ES-504 DII relative to RVP_17D_-WT **(Figure 3D)** and a smaller (∼3–4-fold), but statistically significant, increase in titer with RVP_ES-504_–17D DII relative to RVP_ES-504_-WT **(Figure 3E)**. The DI chimeras afforded smaller, but concordant, changes in neutralization sensitivity, whereas the DIII chimeras were unaffected **(Figure 3D-E)**. These findings indicate that one or more sequence polymorphisms in DII of glycoprotein E largely account for the reduced sensitivity of YFV-ES-504 to neutralization by YF-17D vaccinee sera.

**Figure 3.**
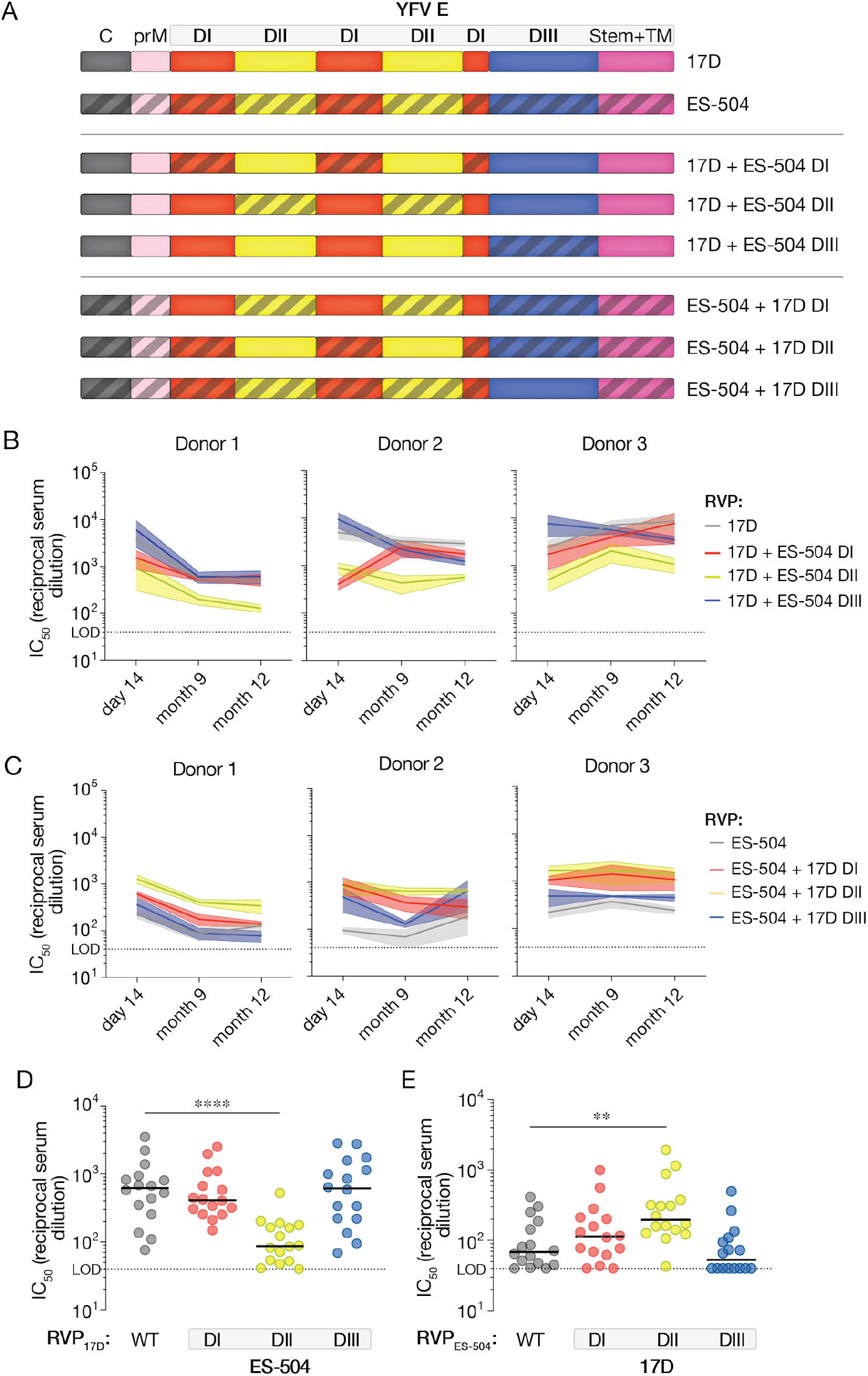
Neutralizing activities of vaccinee sera against RVPs bearing 17D/ES-504 domain-swapped E proteins. (**A**) Schematic of the 17D/ES-504 E protein chimeras. (**B–C**) Serum neutralizing titers for the three U.S. donors (see Figure 1A) at the indicated times post-vaccination against the indicated RVPs. Means±SEM, n=6–12 from 2-4 independent experiments. (**D–E**) Serum neutralizing titers for a U.S. vaccinee cohort (n=16 donors; see Figure 1E) against the indicated RVPs. n=6–12 from 2–4 independent experiments. Parental RVP_17D_ and RVP_ES-504_ controls are from Figure 1E. Groups in D–E were compared by two-way ANOVA followed by Tukey’s correction for multiple comparisons. **, P<0.002. ****, P<0.0001. Only the significant comparisons are shown. In all panels, lines indicate group medians. LOD, limit of detection.

### Two sites in domain II influence YFV neutralization by YF-17D vaccinee sera

A comparison of the amino acid (aa) sequences of African (17D, Asibi, China) and South American YFV strains (ES-504) revealed nonconservative changes unique to YFV-ES-504 at five positions in DII: H67N, A83E, D270E, N271S, and N272K **(Figure 4A)**. The first two positions (Site 1: residues 67 and 83) map within or near the binding footprints of the recently described protective nAb 5A **(Figure 4)** (Lu et al., 2019) and a large cohort of potent, 5A-competing neutralizers we recently isolated from YF-17D vaccinees (Wec et al., 2020). These nAbs recognize overlapping, fusion loop-proximal DII epitopes in E proteins from African strains (Lu et al., 2019; Wec et al., 2020). The last three polymorphic residues (Site 2: residues 270–272) are located in the *kl* loop that forms part of the hinge region connecting DII and DI ((Lu et al., 2019) and PDB: 6EPK) **(Figures 4B–C)**.

**Figure 4.**
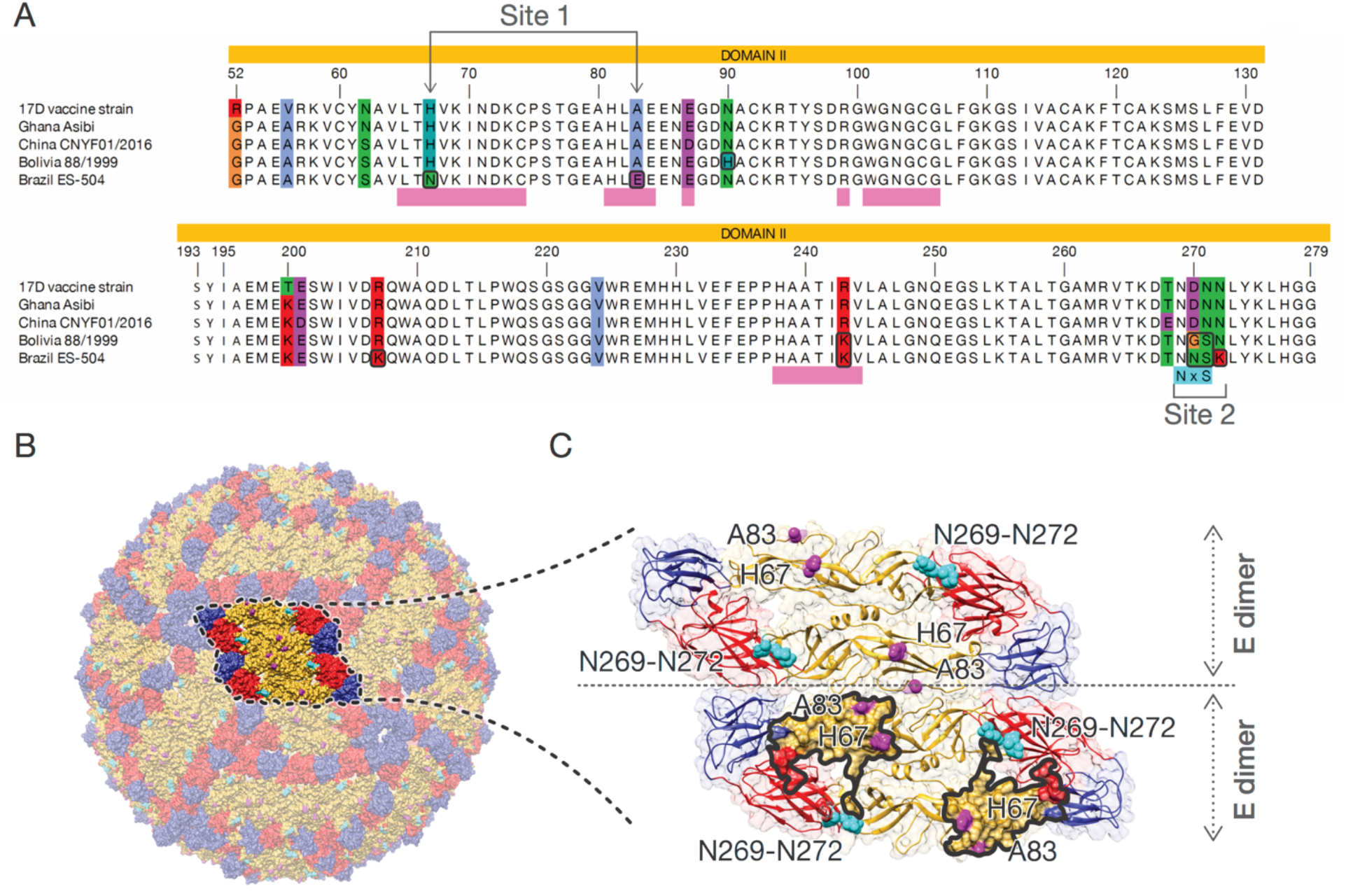
YFV strain-dependent sequence polymorphisms in DII of the E protein. (**A**) Amino acid sequence alignments of the portion of E corresponding to DII are shown for the indicated sequences. The colored residues in the alignment correspond to positions where residue identities are not identical in all five sequences. Polymorphic sites 1 and 2 are indicated below the alignment. (**B**) Amino acid positions with unique residue identities in YFV-ES-504 E are shown on a YFV virion, which was constructed in Chimera (Pettersen et al., 2004) using the atomic coordinates of YFV E (Lu et al., 2019) and the symmetry matrix of the dengue virus 2 virion (Zhang et al., 2004). (**C**) Close-up view displaying two neighboring dimers. Highlighted surface area in the lower dimer corresponds to the nAb 5A binding interface as dictated by PISA (relative buried surface area ≥ 10%) (Krissinel and Henrick, 2007). Residue numbering corresponds to that of the mature YFV-17D E protein.

To investigate the effect of these localized aa substitutions, we evaluated the neutralization sensitivity of RVPs bearing additional 17D-ES-504 E chimeras in which these two sites (Site 1: residues 67 and 83; Site 2: residues 270–272) were exchanged **(Figure 5A)**. Swapping Site 1 between YFV-17D and -ES-504 E proteins reduced the neutralizing activity of the DII-specific vaccinee nAbs, concordant with the preponderance of known 5A-competing nAbs in this panel **(Figures 5B–C and S3)**. Strikingly, however, these Site 1 chimeras had little or no impact on viral neutralization by the YF-17D vaccinee sera **(Figure 5D–E)**. Conversely, whereas the DII nAbs (**Figure 5B–C)** and their 5A-competing subset **(Figure S3)** were much less affected by the Site 2 chimeras, the introduction of ES-504 Site 2 into RVP_17D_ was associated with substantial losses in serum neutralizing activity **(Figure 5D)**. The reciprocal RVP_ES-504_ bearing 17D Site 2 showed a smaller increase in sensitivity to the vaccinee sera **(Figure 5E)**, but combining the Site 1 and Site 2 changes in each strain background afforded larger shifts in serum neutralizing titer approaching those observed with the DII chimeras **(Figures 5D–E)**. Taken together, these findings indicate that sequence polymorphisms at Sites 1 and 2 in DII largely account for the reduced neutralizing activity of YF-17D vaccinee sera against YFV-ES-504, especially when both sets of 17D→ES-504 changes are simultaneously present. They also suggest that this phenotype observed with polyclonal sera cannot be readily attributed to changes in activity of the potently neutralizing, 5A-competing, nAbs alone.

**Figure 5.**
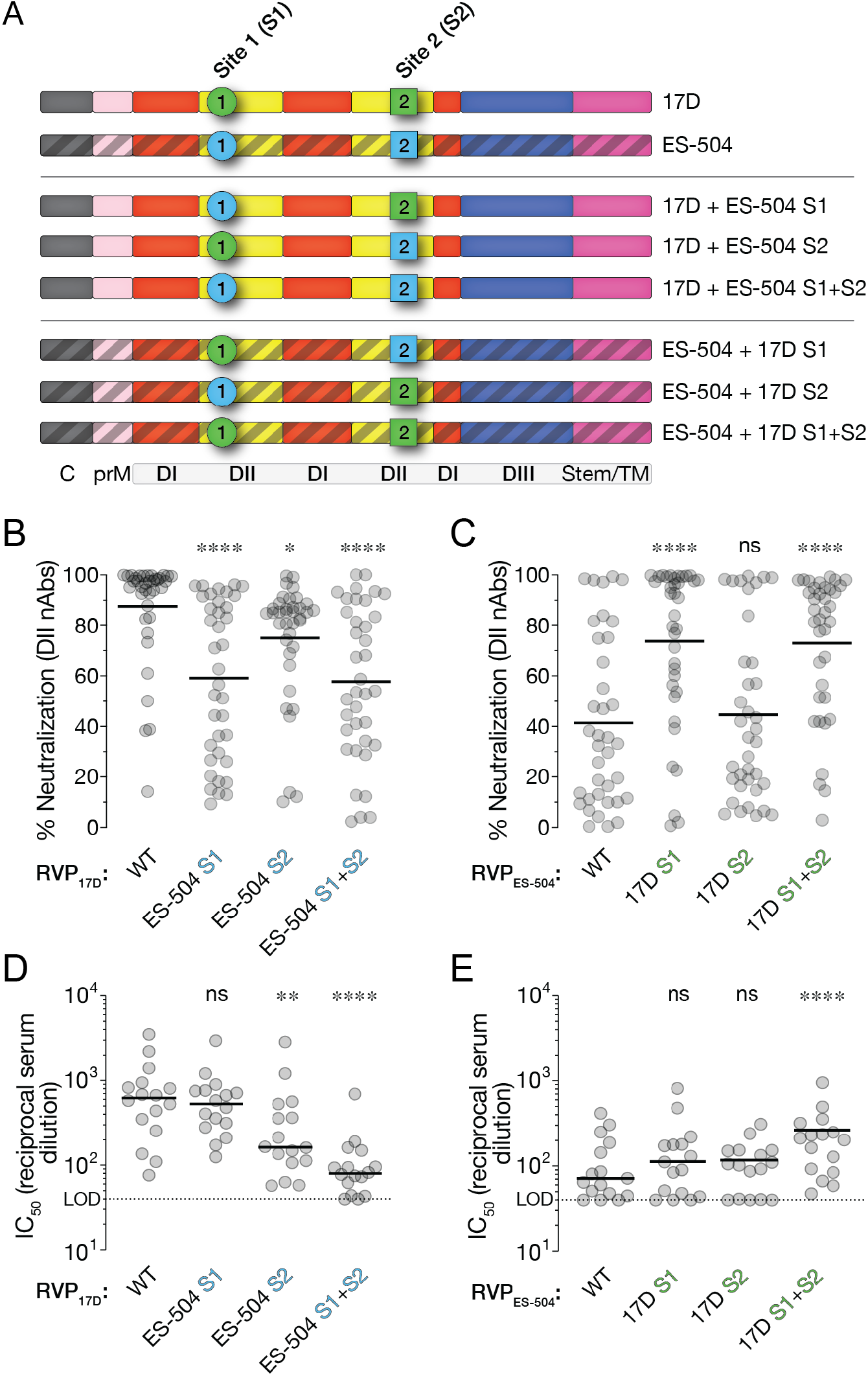
Effects of polymorphisms at Sites 1 and 2 on the neutralizing activities of vaccinee nAbs and sera. (**A**) Schematic of YFV-17D and -ES-504 E proteins chimerized at Sites 1 and 2. (**B–C**) Neutralizing activities of selected DII-specific nAbs (10 nM) against the indicated RVPs. Means, n=6–12 from 2–4 independent experiments, Parental RVP_17D_ and RVP_ES-504_ controls are from Figure 2. (**D–E**) Serum neutralizing titers for a U.S. vaccinee cohort (n=16 donors; see Figure 1E) against the indicated RVPs. Means, n=6–12 from 2–4 independent experiments. Parental RVP_17D_ and RVP_ES-504_ controls are from Figure 1E. Groups were compared by two-way ANOVA followed by Tukey’s correction for multiple comparisons. **, P<0.002. ****, P<0.0001. ns, not significant. In all panels, lines indicate group medians. LOD, limit of detection.

### *N*–glycosylation at residue 269 and Asn-to-Lys change at residue 272 both contribute to YFV-ES-504 neutralization resistance

The central portion of the *kl* loop contains Site 2 ((Lu et al., 2019) and PDB: 6EPK) and is not known to be a critical binding determinant of YFV E-specific mAbs (Daffis et al., 2005; Lu et al., 2019). Through both manual inspection and a search for eukaryotic linear motifs, we identified a putative *N*–linked glycosylation site (NXS/T sequon) at Site 2 residues 269–271 (NNS) in YFV-ES-504 but not YFV-17D or -Asibi (NDN) **(Figure 4A)**. To obtain experimental evidence for viral strain-dependent E glycosylation at this position, we subjected recombinant, soluble YFV-Asibi and YFV-ES-504 E proteins (sE) produced in S2 cells to mass spectrometry. Consistent with the bioinformatic predictions, LC-MS/MS analysis of tryptic glycopeptides revealed a high degree of *N*–glycan occupancy at aa residue Asn 269 in YFV-ES-504 sE (>90%) but no detectable N– glycosylation at the same position in YFV-Asibi sE **(Figure 6A)**.

**Figure 6.**
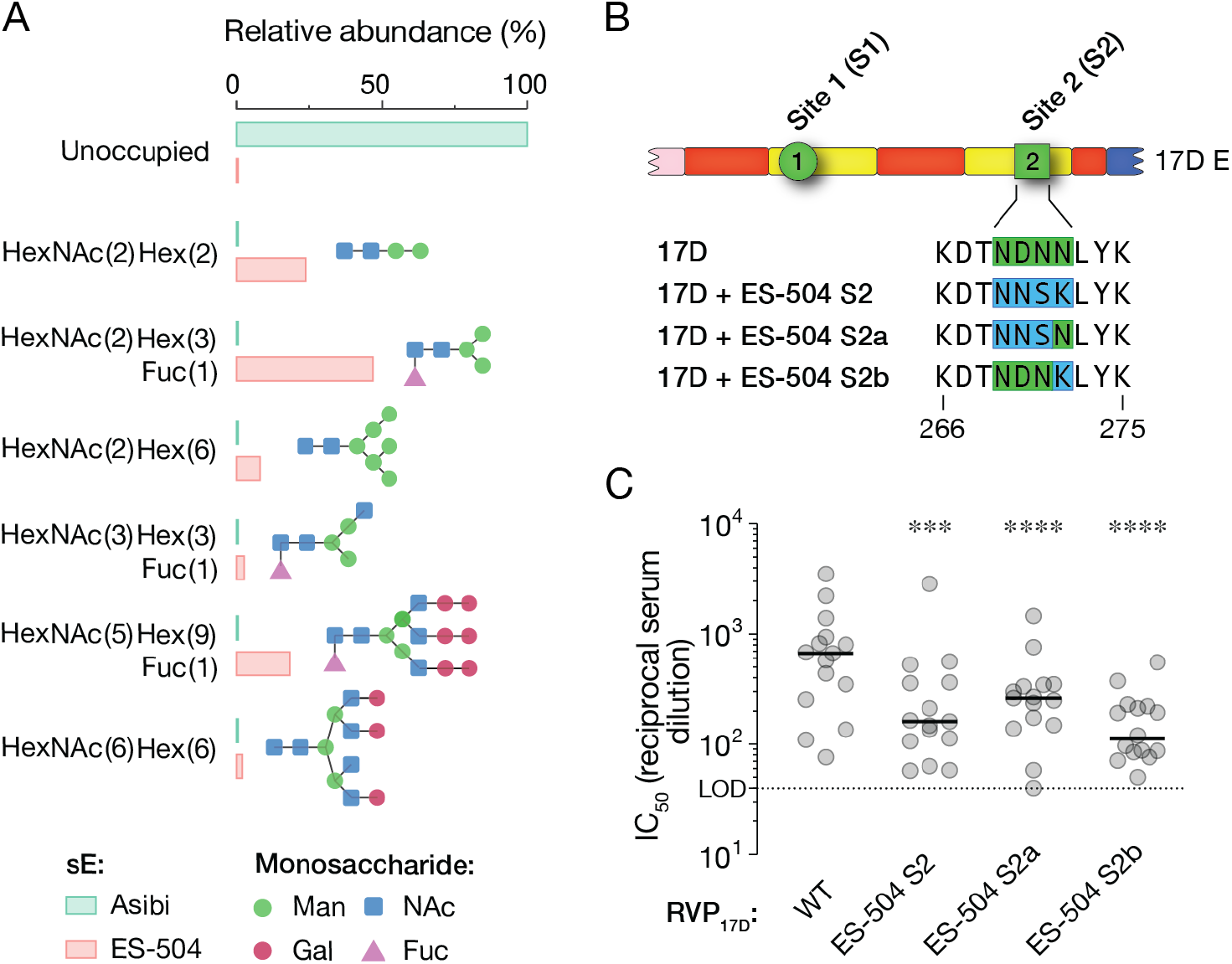
Effects of polymorphisms at subsite 2a and 2b on *N–*linked glycosylation of YFV E and neutralization sensitivity to vaccinee sera. (**A**) Occupancy and relative abundance of the indicated glycan structures at residue Asn 269 in recombinant soluble E proteins (sE) from YFV-Asibi and YFV-ES-504 E were determined by mass spectrometry. (**B**) Schematic of YFV-17D E protein chimerized at subsites 2a and 2b. (**C**) Serum neutralizing titers for a U.S. vaccinee cohort, n=15, against the indicated RVPs. Means, n=6 from 2 independent experiments. Parental RVP_17D_ controls are from Figure 1E. Groups were compared by two-way ANOVA followed by Tukey’s correction for multiple comparisons. ***, P<0.0002. ****, P<0.0001. In all panels, lines indicate group medians. LOD, limit of detection.

To independently assess the roles of this sequon (Site 2a) and the Asn-to-Lys change at residue 272 (N272K; Site 2b) in determining the Site 2 neutralization phenotype, we tested two additional RVP_17D_ chimeras bearing ES-504 Site 2a or Site 2b **(Figure 6B)**. Changes at both subsites reduced the neutralizing activity of YF-17D vaccinee sera, with the Site 2b change having the larger effect **(Figure 6C)**.

### Changes at Sites 1 and 2 show distinct patterns of conservation among South American E sequences

We aligned a representative set of 82 non-redundant YFV E protein sequences encompassing the sequence space of 281 fully sequenced YFV E genes (see Methods) **(Supplementary Excel File 2)**. This alignment revealed that the Site 1 and 2 changes were unique to YFV sequences of South American origin **(Figure 7)** but also uncovered subclade-specific differences in the occurrence of these polymorphisms. Specifically, the 17D→ES-504 changes at Site 1 and Site 2b were observed in nearly all sequences from the subclade to which YFV-ES-504 belongs (South American ‘genotype I’), but not in any of the unique South American ‘genotype II’ sequences available for analysis. By contrast, both genotype I and II strains encoded a sequon at Asn 269, albeit with different sequences (genotype I, NNS; genotype II, NGS) **(Figure 4A)**. RVPs bearing E from a prototypic genotype II strain, YFV-Bolivia (YFV/BOL/88/1999), also displayed resistance to neutralization by YF-17D vaccinee sera **(Figure S4)**, although at reduced levels relative to YFV-ES-504. YFV-Bolivia’s intermediate phenotype may reflect that it and other South American genotype II strains combine *N–*glycosylation at Asn 269, a YFV-ES-504–like signature, with YFV-17D/Asibi-like signatures at Sites 1 and 2b **(Figures 6B and 4A)**, although other sequences polymorphic between YFV-ES-504 and YFV-Bolivia may also play a role. Taken together, our findings indicate that genetic variation at two distinct sites in DII influences the reduced sensitivity of YFV-ES-504 and other South American strains to neutralization by YF-17D vaccinee sera.

**Figure 7.**
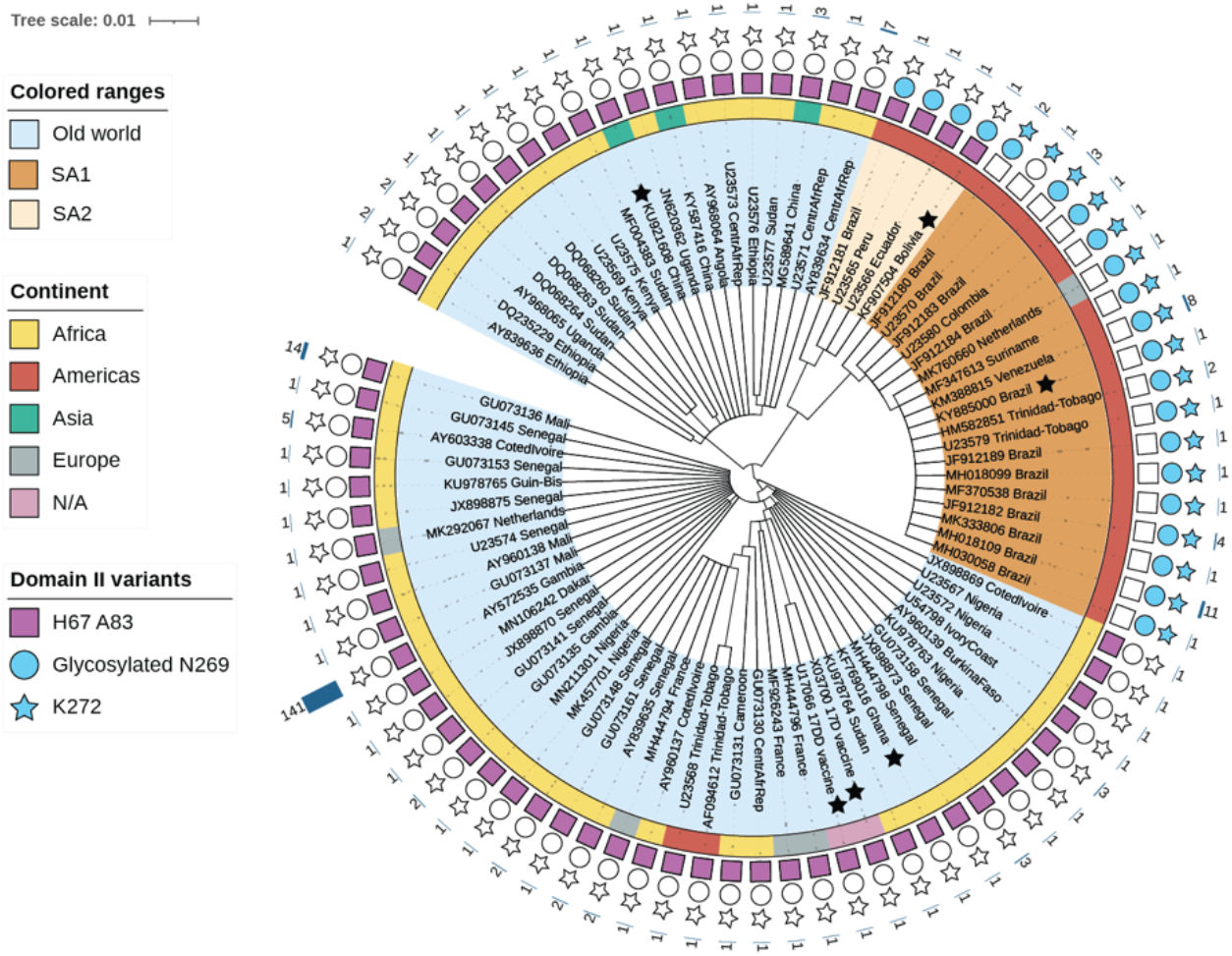
Genetic makeup of YFV strains at polymorphic sites 1 and 2. Amino acid-based tree depicting E proteins from representative YFV strains (scale bar represents the number of amino acid substitutions per site). Colored ranges denote viral origin and clade(s): i) Old World (light blue), ii) SA1 (South American genotype 1, brown). SA2 (South American genotype 2, beige). Black stars indicate YFV sequences used in this study. The inner ring is color-coded according to the continent of origin. Note that all sequences of apparent Asian or European origin are ex-African (if clustering with Old World) or ex-South American (if clustering with SA1). N/A, not applicable. Sequence features are represented with a square (Site 1: H67 A83), circle (Site 2a: predicted glycan at N269), or star (Site 2b: K272). Features present in sequence are represented with filled shapes. The outer char bar denotes the number of strains being represented at each branch of the tree.

## Discussion

Mass vaccination campaigns have historically been, and remain, a vital tool in the public health armamentarium against YFV. Despite considerable efforts, however, attaining sufficient population vaccine coverage in endemic areas and ‘surging’ vaccination during epidemics has proven challenging. In Brazil, shortages in the locally manufactured and highly effective YFV-17D vaccine represent one major logistical hurdle to widespread vaccination and necessitated the use of dose-sparing strategies to maximize coverage during the 2016–19 epidemic (Boëte, 2016; de Oliveira Figueiredo et al., 2020). A further challenge to maintaining long-term population immunity is the World Health Organization’s recommendation that individuals receive a single lifetime dose of the vaccine. The emerging consensus points instead to the need for a second dose to boost waning immunity and mitigate the demonstrated risk of re-infection in YF-vaccinated individuals (Campi-Azevedo et al., 2019b, 2019a; Estofolete and Nogueira, 2018; Vasconcelos, 2018).

Measurement of antiviral neutralizing antibodies in human sera is the most widely used biomarker for YFV vaccine immunogenicity and provides a key correlate of vaccine protection (Mason et al., 1973; Plotkin, 2010). Here, we identify a previously unrecognized factor — the choice of indicator viral strain — that further complicates the interpretation of serum neutralizing titers in South American vaccinees. Specifically, YFV neutralizing titers in vaccinee sera are conventionally determined using the YF-17D vaccine strain or one of its variants (e.g., YF-17DD), ultimately derived from the virulent African strain, YFV-Asibi. We found that neutralizing antibody responses induced by YF-17D vaccination were markedly attenuated against South American strains YFV-ES-504 and YFV-Bolivia (belonging to genotypes I and II, respectively) **(Figures 1 and S4)**. Because the genetic features of these sequences that influence neutralization sensitivity appear to be shared by most South American YFV strains (see below), we conclude that available serological data likely overestimate the potency and longevity of neutralizing antibody responses in South American vaccinees.

We genetically mapped the reduced neutralization sensitivity of YFV-ES-504 versus YFV-17D to the central domain II (DII) of the E protein **(Figure 4)**, which is the major target of polyclonal neutralizing antibody responses in flavivirus vaccinees and convalescents (Vratskikh et al., 2013; Wec et al., 2020). Further genetic dissection of DII revealed two polymorphic sites. The first (Site 1) is a fusion loop-proximal sequence containing or overlapping the epitopes of the protective nAb, 5A (Lu et al., 2019) **(Figure 4C)** and those of a large panel of 5A-competing nAbs we recently isolated from YF-17D vaccinees (Wec et al., 2020) **(Figures 2 and S3 and Supplementary Excel File 1)**. Site 2 is a sequence at the apex of the *kl* loop, which forms part of the hinge region between DI and II **(Figure S5A–B)**. We found that the 5A-competing, DII-specific nAbs were much more sensitive to the changes at Site 1 than to those at Site 2 **(Figures 5B-C and S3)**, whereas YF-17D vaccinee sera displayed the opposite pattern, especially in the YFV-17D genetic background **(Figure 5D-E)**. Nevertheless, combining the Site 1 and Site 2 polymorphisms afforded the largest changes in YFV neutralization by vaccinee sera **(Figure 5D–E)**. These findings suggest that YF-17D vaccination elicits both currently undefined mAb specificities with key roles in viral neutralization.

We provide genetic and biochemical evidence for two determinative changes at Site 2 associated with YFV-17D→ES-504 neutralization resistance: the acquisition of an *N*–linked glycan at Asn 269 and an Asn-to-Lys change at residue 272 (N272K) **(Figure 6)**. Although *N–* glycosylation of YFV E has been sporadically observed (e.g., at Asn 153, seen in the YFV-17DD vaccine variant only (Post et al., 1992) **(Supplementary Excel File 2)**, this protein is generally reported to not bear any *N–*glycans, possibly because only African clade sequences have been used in most prior analyses (Carbaugh and Lazear, 2020; Post et al., 1992). Indeed, we could find only one report in which the presence of a sequon at Asn 269 was noted in five South American YFV sequences (Chang et al., 1995). Here, we show that predicted *N*–glycosylation at Asn 269 is almost universally observed in South American YFV strains, including both genotypes I and II and in both ‘Old’ and ‘New’ genotype I strains (Carrington and Auguste, 2013; Gómez et al., 2018), but not in any African or other Old World YFV strains sequenced to date **(Figure 7)**. Intriguingly, South American genotype I and II strains possess distinct Asn 269 sequons (NNS and NGS, respectively), raising the possibility that they may have independently evolved this feature. The N272K change at the DI-DII hinge in the YFV E protein has not been previously described, to our knowledge.

An extensive body of work has shed light on the patterns and functional significance of *N–*linked glycosylation in flavivirus E proteins (Carbaugh and Lazear, 2020; Yap et al., 2017). Although the number of *N*–glycans vary in a virus-, strain-, and lineage-dependent manner, recurrent *N*–glycosylation is only observed at two positions: Asn 67 in DII and Asn 153/154 in the variable ‘150-loop’ sequence in DI. Current evidence indicates that *N–*linked glycans at Asn 67 and/or Asn 154 play cell type- and host-dependent roles in flavivirus multiplication, by virtue of which they can impact virulence, pathogenesis, and mosquito-borne transmission (Carbaugh and Lazear, 2020; Yap et al., 2017). To our knowledge, *N–*glycosylation at Asn 269 in the DI-DII hinge has not been observed in any other flavivirus E protein to date. We speculate that acquisition of the Asn 269 glycan by South American YFV E proteins was an early adaptation event that afforded the virus a selection advantage in its new sylvatic environments in the New World.

Our observation that two distinct biochemical changes at the DI–DII hinge in the *kl* loop could independently impact viral neutralization sensitivity by polyclonal sera raises the possibility that they induce allosteric changes in the E protein and influence the structural dynamics of the entire glycoprotein shell (“breathing”) in a manner that regulates the availability of other epitopes, as has been described for other flaviviruses (Barba-Spaeth et al., 2016; Dejnirattisai et al., 2015; Dowd et al., 2015; Goo et al., 2017, 2018). Indeed, the asymmetric effects of the YFV-17D/ES-504 Site 2 chimeras on serum neutralization **(Figure 5C–D)** appear consistent with such allosteric models. The analysis of predicted and known B-cell epitopes in YFV E and its structural superposition with other flavivirus E:nAb complexes identified multiple nAb footprints in proximity to residues in the *kl* loop **(Figure S5C–L and Supplementary Excel File 3)**. Several of these nAbs recognize complex quaternary epitopes that span multiple E subunits in the intact viral particles of other flaviviruses (reviewed in (Dussupt et al., 2020; Sevvana and Kuhn, 2020). Indeed, previous work with both YFV and DENV suggests that quaternary epitope binders may constitute a substantial fraction of the neutralizing activity in human immune sera (de Alwis et al., 2012; Vratskikh et al., 2013). We anticipate that such antibodies are underrepresented in our nAb panel, which was isolated by B-cell sorting with a largely monomeric recombinant sE protein (Wec et al., 2020). We propose that they comprise part of the large reservoir of neutralizing activity in YF-17D vaccinee sera that is affected by the polymorphisms at Site 2 but not Site 1.

Although the genetic distinctiveness of the South American and African YFV clades has been previously noted, the implications of specific clade- and lineage-dependent differences for viral transmission, virulence, and immune evasion have not been investigated. Here, for the first time, we identify conserved genetic and biochemical features unique to South American YFV strains that influence their reduced susceptibility to nAbs in human vaccinee sera. These results argue for a reevaluation of current approaches to YFV immunological surveillance in South America. They also point to an urgent need to expand and update our understanding of the quantitative relationship(s) among vaccine-induced neutralizing antibodies, T cells, and vaccine-mediated protection against YFV, especially in light of the genotype-dependent effects uncovered herein. Finally, our findings provide a roadmap to explore updates to the current gold-standard vaccine that forms the basis of WHO’s comprehensive global strategy to eliminate YF epidemics (EYE) (WHO, 2018).

## Acknowledgments

We thank I. Gutierrez, E. Valencia and L. Polanco for laboratory management and technical support. We thank Dr. Ted Pierson for his provision of the WNV-based RVP system. This work was funded by the U.S. National Institutes of Health grant R01AI132633 (to K.C.), “Preventing and Combating the Zika Virus” MCTIC/FNDCT-CNPq/MEC-CAPES/MS-Decit. (Grant no. 440865/2016-6) and Conselho Nacional de Desenvolvimento Científico e Tecnológico (Grant no. 312446/2018) awarded to M.C.B. M.C.B. is a recipient of CNPq fellowship for Productivity in Technological Development and Innovative Extension (Grant 309471/2016-8). Si.Si. and St.St. were supported by the Leukemia Research Foundation (Hollis Brownstein New Investigator Research Grant), AFAR (Sagol Network GerOmics award), Deerfield (Xseed award) and the NIH Office of the Director (1S10OD030286-01). Y.S. gratefully acknowledges the NIH grant R01GM129350-04. C.A.C. is a recipient of the PhD scholarship provided by the Coordenação de Aperfeiçoamento de Pessoal de Nível Superior - Brasil (CAPES)

## Author Contributions

Conceptualization, D.H., G.L., A.Z.W., L.M.W., M.C.B., K.C.; Investigation, D.H., G.L., A.Z.W, N.D.F., L.M.S.R, Y.S., St.St., N.P.L., A.T.; Formal analysis, D.H., K.C.; Resources, C.A.C, C.L.M., M.S., V.I.A., Z.A.B., E.G K, L.M.W., M.C.B., K.C.; Data Curation, D.H., G.L., A.Z.W., M.S.; Writing - Original Draft, D.H., G.L., A.Z.W., K.C; Writing - Review & Editing: All authors; Visualization, D.H., G.L., A.Z.W., K.C.; Supervision, Z.A.B., D.I.K., E.K., Si.Si., L.M.W., M.C.B., K.C.; Project Administration, D.H., K.C.; Funding Acquisition, E.G.K., M.C.B., K.C.

## Declaration of Interest

K.C. is a member of the scientific advisory boards of Integrum Scientific, LLC and Biovaxys Technology Corp. A.Z.W., M.S., and L.M.W. are/were employees of Adimab, LLC, and may hold shares in Adimab, LLC. L.M.W. is an employee of Adagio Therapeutics, Inc., and holds shares in Adagio Therapeutics, Inc. C.L.M and Z.A.B are employees of Mapp Biopharmaceutical, Inc.

## STAR Methods

### RESOURCE AVAILABILITY

#### Lead Contact

Further information and requests for resources and reagents should be directed to and will be fulfilled by the Lead Contact, Kartik Chandran.

#### Materials Availability

Plasmids in this study, together with documenting information, will be made available upon request and following execution of a UBMTA between Albert Einstein College of Medicine and the recipient institution. Primary data are available on request. This study did not generate any new software code.

### EXPERIMENTAL MODEL AND SUBJECT DETAILS

#### The U.S. cohort

Informed consent to participate in this study was obtained before vaccination. Study subjects (donors 1 to 3) were vaccinated with the YFV-17D Stamaril vaccine. Heparinized blood (50 to 100 cc) was obtained from subjects before vaccination and for over one year following vaccination as described previously (Wec et al., 2020). In addition, from donors 2 and 3 after 3 and 2 years respectively. Additional consented volunteers (donors 9 to 18) with history of YFV vaccination (Stamaril or YF-Vax, both YF-17D-204 virus strain) were recalled for plasma collection at the indicated time points ranging up to 20 years post first YFV vaccination. Samples were processed in the Immune Monitoring and Flow Cytometry Core laboratory at the Geisel School of Medicine at Dartmouth College to obtain plasma and to isolate peripheral blood-derived B cells. Isolated cells and plasma were stored frozen in aliquots at −80 °C. This study complies with all relevant ethical regulations for work with human participants and was approved by the Committee for the Protection of Human Subjects, Dartmouth-Hitchcock Medical Center, and Dartmouth College. Additional blood samples were collected from healthy adult volunteers after giving informed consent (donors 19 to 22) with history of YFV vaccination (Stamaril or YF-Vax). Blood was drawn by venipuncture to collect serum. Serum was centrifuged, aliquoted and stored at −80°C. The study protocol was approved by the Institutional Review Board (IRB) of the Albert Einstein College of Medicine (IRB number 2016-6137).

#### The Brazilian cohort

Study subjects (donors 4 to 8 and donors 23 to 46) were selected between October 2011 and April 2014, among people who attended the Reference Centre of Special Immunobiologicals at the Clinics Hospital in the University of São Paulo (CRIE-HCFMUSP). Individuals looking for the attenuated yellow fever vaccine, either HIV-positive or not, were evaluated by medical personnel and invited to participate in the study, if they were naïve for the vaccine. They were between 18 and 59 years old and should not have auto-immune diseases or other health issues that could impact on the vaccination. A team researcher clarified the terms of the study, presenting the aims of the research, the potential risks, and the procedures. The individuals who agreed to participate signed an informed consent form. The study subjects were then vaccinated with the YF-17DD attenuated vaccine (Biomanguinhos, Fiocruz), and blood was drawn from 5 donors (donors 4 to 8) over 1 year. Another 24 donors (donor 23 to 46) had their first vaccine dose at the same center and samples were collected one to two years post-vaccination. Whole blood from vaccinees was collected in EDTA tubes. For plasma separation samples were centrifuged at 1800 rpm for 10 minutes. The plasma was collected and centrifuged again at 2800 rpm for 10 minutes to remove cellular debris. After centrifugation, plasma was aliquoted and stored at −20°C for further use. The research was approved by the Ethics Committee for Research Project Analysis (CAPPesq) from the Clinics Hospital at the University of São Paulo (CAAE: 83631318.000.068).

#### Cells

Human hepatoma-derived Huh 7.5.1 cells (originally from Dr. Frank Chisari and received from Dr. Jan Carette) were passaged using 0.05% Trypsin/EDTA solution (Gibco) every three to four days and maintained in Dulbecco’s Modified Eagle Medium (DMEM high glucose, Gibco) complemented with 10% fetal bovine serum (FBS, heat-inactivated, Atlanta Biologicals), 1% Penicillin/Streptomycin (P/S, Gibco), 1% Gluta-MAX (Gibco) and 25 mM HEPES (Gibco).

Human embryonal kidney 293FT cells, purchased from Thermo Fisher, were passage every 3 to 4 days using 0.05% Trypsin/EDTA solution (Gibco) and maintained in DMEM high glucose (Gibco) complemented with 10% FBS (heat-inactivated, Atlanta Biologicals), 1% P/S (Gibco), 1% Gluta-MAX (Gibco) and 25 mM HEPES (Gibco).

Vero cells (ATCC-CCL81) were maintained in Earle’s 199 medium (Gibco) supplemented with 5% fetal bovine serum (FBS; Gibco), 0.25% sodium bicarbonate (Sigma-Aldrich), and 40 mg/mL gentamicin (Gibco) in an incubator at 37°C with wet atmosphere and 5% CO2.

C6/36 cells, kindly provided by Dr. Anna-Bella Failloux (Institut Pasteur, Paris), were grown in Leibovitz’s L-15 medium (Gibco), supplemented with 5% FBS, 10% tryptose broth, and 40 mg/mL gentamicin in an incubator at 28°C.

#### Virus generation

Yellow Fever virus 17D (YFV-17D) was obtained from BEI Resources (Cat# NR-115). Huh 7.5.1 cells were seeded into 15cm dishes and confluent cells were infected using 90 µL of passage 2 stock of the YFV-17D supernatant in 3 mL infection media (DMEM low glucose (Gibco), 7% FBS (heat-inactivated, Atlanta Biologicals), 1% P/S (Gibco), 1% Gluta-MAX (Gibco), 25 mM HEPES (Gibco)) for 1 hour at 37 °C and 5% CO2. Three days later supernatant was harvested and cleared from cell debris by centrifugation twice at 4,000 rpm for 15 min at 4 °C. The cleared viral supernatant was aliquoted and stored at −80 °C.

YFV ES-504 was isolated from the blood of a non-human primate found dead in Espírito Santo, Brazil, in February 2017 (Bonaldo et al., 2017). The viral stock was produced in C6/36 cells. Cells were seeded at a density of 80,000 cells/cm² in a T-75 flask 24 hours before infection with 3 mL of viral isolate suspension. After 1 hr incubation at 28°C, 40 mL of supplemented Leibovitz’s L-15 medium was added. After 8 days, the supernatant was centrifuged at 700 x g for 10 min at 4 °C to remove cell debris, aliquoted, and stored at −80°C.

YFV-17DD was recovered from the lyophilized vaccine ampoule produced by Biomanguinhos-FIOCRUZ/RJ with supplemented Earle’s 199 medium. The viral stock was obtained in Vero cells seeded at 62,500 cells/cm² in a T-150 culture flask. Viral adsorption was carried out for 1 hr, after which 40 mL of medium was added. Cells were observed daily until the cytopathic effect was detectable. The cell supernatant was centrifuged at 700 x g for 10 min at 4 °C to remove cell debris, aliquoted, and stored at −80°C.

#### Reporter virus particle (RVP) generation

West Nile virus (WNV) subgenomic replicon-expressing plasmid pcDNA6.2-WNIIrep-GFP/zeo was a generous gift from Ted Pierson (Pierson et al., 2006). The structural proteins C, prM and E of YFV-17D, ES-504, China and Bolivia (Genbank: X03700.1, KY885000.2, KU921608.1 and KF907504.1 respectively) as well as their mutated forms were synthesized and codon-optimized for expression in human cells by Epoch Life Science, Inc. and subcloned into the pCAGGS vector for mammalian expression. For RVP_YFV-17D/ES-504_ chimeras, DI (amino acid residues 1–51, 132–192 and 280–295), DII (residues 52–131 and 193–279) and DIII (residues 296–394) were exchanged. For RVP_17D_ or RVP_ES-504_ carrying site mutations, Site 1 (residues 67 and 83) and Site 2 (residues 270–272 DNN/NSK were interchanged (Figure 4). For RVP_17D_ NSN/NDK mutations at 270–272 were introduced accordingly (Figure 6). AA labelling according to the E protein sequence of YFV-17D.

293FT cells (Thermo Fisher) were transfected with 12 µg of total DNA per 15 cm plate in a ratio of 1:3 using the WNV replicon plasmid and the respective virus C-prM-E plasmid. Eight hours post-transfection, media was exchanged to low glucose DMEM (Gibco) media containing 5% FBS (heat-inactivated, Atlanta Biologicals) and 25mM HEPES (Gibco). After 3 to 4 days at 37 °C under 5% CO2, the cell supernatant containing RVPs was harvested and to remove the cell debris centrifuged twice for 15 minutes at 4,000 rpm at 4 °C. The viral supernatant was pelleted using a SW28 rotor (Beckman Coulter) in a Beckman Coulter Optima LE-80K ultracentrifuge for 4 hours at 4 °C through a 2 mL cushion of 30% (v/v) D-sucrose in PBS (pH 7.4). The pellet was resuspended overnight on ice in 100 µl PBS (pH 7.4), aliquoted and stored at −80°C till further use.

#### Production of recombinant YFV antigens

The coding sequence for the entire prM and soluble E (sE) regions (amino acid residues 122–678 of the YFV polyprotein) of the YFV Asibi Strain (Uniprot ID: Q6DV88) or YFV ES-504 strain (Uniprot ID: A0A1W6I1A1) were cloned into an insect expression vector encoding a C-terminal double strep tag, pMT-puro. The expression construct design was based on previously published structures of flavivirus antigens (Modis et al., 2003, 2005; Rey et al., 1995). The YFV prM/E constructs were used to generate an inducible, stable Drosophila S2 line. Protein expression was induced with addition of copper sulfate and allowed to proceed for 5 to 7 days. Recombinant proteins were affinity purified from the culture supernatant with a StrepTrap HP column (GE Healthcare) and an additional purification step was carried out using size-exclusion chromatography step using an S200 Increase column (GE Healthcare). The final protein preparations were stored in PBS (pH 7.4) supplemented with an additional 150 mM NaCl, aliquoted and stored at −70 °C.

#### Monoclonal antibodies

The isolation and characterization of monoclonal antibodies used in this study have been described previously in detail in (Wec et al., 2020). In short, IgGs were expressed in S. cerevisiae cultures grown in 24-well plates, as described previously (Bornholdt et al., 2016). After 6 days, the cultures were harvested by centrifugation and IgGs were purified by protein A-affinity chromatography. The bound antibodies were eluted with 200 mM acetic acid/50 mM NaCl (pH 3.5) into 1/8th volume 2 M Hepes (pH 8.0) and buffer-exchanged into PBS (pH 7.0).

### METHOD DETAILS

#### BioLayer Interferometry Kinetic Measurements (BLI)

For monovalent apparent KD determination, IgG binding to recombinant YFV E antigen was measured by biolayer interferometry (BLI) using a FortéBio Octet HTX instrument (Molecular Devices). The IgGs were captured (1.5 nm) to anti-human IgG capture (AHC) biosensors Molecular Devices) and allowed to stand in PBSF (PBS with 0.1% w/v BSA) for a minimum of 30 min. After a short (60 s) baseline step in PBSF, the IgG-loaded biosensor tips were exposed (180 s, 1000 rpm of orbital shaking) to YFV E antigen (100 nM in PBSF) and then dipped (180 s, 1000 rpm of orbital shaking) into PBSF to measure any dissociation of the antigen from the biosensor tip surface. Data for which binding responses were > 0.1 nm were aligned, inter-step corrected (to the association step) and fit to a 1:1 binding model using the FortéBio Data Analysis Software, version 11.1.

#### Microtiter neutralization assays

Plasmas were serially diluted in DMEM high glucose medium (Gibco) containing 10% heat-inactivated FBS (Atlanta Biologicals), 1% Gluta-MAX (Gibco), 1% P/S (Gibco) and 25 mM HEPES (Gibco). Dilutions were incubated at room temperature (RT) with YFV-17D or RVPs for one hour. Monoclonal antibodies at a concentration of 10nM were pre-incubated with RVPs for 1 hour at RT before adding to cell monolayers. The antibody-virus mixture was added in triplicates to 96-well plates (Costar, cat# 3595) containing 5,000 Huh 7.5.1 cell monolayers seeded the day before and incubated for 2 days at 37°C and 5% CO2. Afterwards, cells were fixed withI 4% paraformaldehyde (Sigma) for 10 mins washed with PBS (pH 7.6), three times. Fixed YFV-17D infected-cells were incubated with a pan-flavivirus mouse mAb 4G2 (ATCC) at 2 µg/ml in PBS containing 3% nonfat dry milk powder (BioRad), 0.5% Triton X-100 (MP Biomedicals), and 0.05% Tween 20 (Fisher Scientific) for one hour at RT. Afterwards, cells were washed and incubated with the secondary antibody conjugated to Alexa Fluor 488 goat anti-mouse (Invitrogen) at a 1:500 dilution for one hour at RT. Finally, YFV-17D and RVP infected cells were washed and nuclei were stained with Hoechst-33342 (Invitrogen) in a 1:2,000 dilution in PBS. Viral infectivity was measured by automated enumeration of Alexa Fluor 488-positive or GFP-positive cells from captured images using a Cytation-5 automated fluorescence microscope (BioTek) and analyzed using the Gen5 data analysis software (BioTek). The half maximal inhibitory concentration (IC50) of the plasma was calculated using a nonlinear regression analysis with GraphPad Prism software. Viral neutralization data were subjected to nonlinear regression analysis to extract the half maximal inhibitory concentration (IC50) values (4-parameter, variable slope sigmoidal dose-response equation; GraphPad Prism).

#### Focus Reduction neutralization (FRNT) assays

Virus-specific neutralizing responses were titrated as previously described in (Magnani et al., 2017). Briefly, sera or antibodies were serially diluted in Earle’s 199 medium (Sigma-Aldrich) supplemented with 5% FBS and incubated 1 h at 37°C with YFV-17DD or -ES504. After incubation, the antibody-virus or sera-antibody mix was added to 96-well plates in triplicates containing Vero CL81 cells monolayers. Plates were incubated for 1.5 h at 37°C. The inocula were discarded, and the cells were overlaid with viral growth medium containing 1% CMC (carboxymethyl cellulose). Plates were incubated at 37°C, 5% CO_2_ for 2-3 days after which cells were fixed with BD Cytofix/Cytoperm solution (BD Biosciences) at room temperature for 30 min, washed twice with PBS, and treated with CytoPerm Wash (BD Biosciences) for 5 min, followed by 1 h incubation with the primary antibody 4G2 (MAB10216, EMD Millipore) diluted 1:2,000. Afterwards, plates were washed three times, followed by an hour-long incubation with a secondary antibody goat anti-mouse IgG conjugated to peroxidase (KPL) or anti-mouse horseradish peroxidase (HRP)-conjugated secondary antibody (115035146, Jackson ImmunoResearch Laboratories). Detection proceeded with the addition of True-Blue Peroxidase Substrate (KPL), following the manufacturer’s instructions. The number of foci was analyzed with a CTL Immunospot instrument. The endpoint titer was determined to be the highest dilution with a 50% reduction (IC_50_) in the number of plaques compared to control wells.

#### Mass spectrometry of glycopeptides

YFV**-**ES-504 and YFV-Asibi proteins (20 µg) were reduced for 1 hour in 200 µl of a buffer containing 8 M urea and 5 mM DTT and were alkylated with 20mM iodoacetamide in the dark for 30 minutes. Next, the protein solution was added to Vivaspin columns (10 kDa) and washed three times with 100 mM ammonium bicarbonate, pH 8.0, for buffer exchange. Finally, 1 µg of sequencing grade trypsin (Promega) diluted in 50 mM ammonium bicarbonate was added to the samples and the mixture was incubated for 18 h at 37°C. After incubation, the peptide solutions were dried in a vacuum centrifuge. Prior to mass spectrometry analysis, samples were desalted using a 96-well plate filter (Orochem) packed with 1 mg of Oasis HLB C-18 resin (Waters). Briefly, the samples were resuspended in 100 µl of 0.1% TFA and loaded onto the HLB resin, which was previously equilibrated using 100 µl of the same buffer. After washing with 100 µl of 0.1% TFA, the samples were eluted with a buffer containing 70 µl of 60% acetonitrile and 0.1% TFA and then dried in a vacuum centrifuge. Samples were resuspended in 10 µl of 0.1% TFA and loaded onto a Dionex RSLC Ultimate 300 (Thermo Scientific), coupled online with an Orbitrap Fusion Lumos (Thermo Scientific). Chromatographic separation was performed with a two-column system, consisting of a C-18 trap cartridge (300 µm ID, 5 mm length) and a picofrit analytical column (75 µm ID, 25 cm length) packed in-house with reversed-phase Repro-Sil Pur C18-AQ 3 µm resin. Peptides were separated using a 135 min linear gradient consisting of 0-32% acetonitrile in 0.1% formic acid over 120 minutes followed by 5 minutes of 32-80% acetonitrile and 5 minutes of 80% acetonitrile in 0.1% formic acid at a flow rate of 300 nl/min. The mass spectrometer was set to acquire spectra in a data-dependent acquisition (DDA) mode. Briefly, the full MS scan was set to 400-2000 m/z in the orbitrap with a resolution of 120,000 (at 200 m/z) and an AGC target of 4×10e5. MS/MS was performed in the orbitrap using an AGC target of 5×10e4 and an HCD collision energy of 30 for fragmentation of glycopeptide ions. Protein Metrics software (Version 3.10) was used to extract the glycopeptide fragmentation data from raw files. N-glycan library from Protein Metrics was used to identify the glycans, together with glycopeptide fragmentation data, which contains b, y and oxonium ions. Glycopeptide with its corresponding extracted ion chromatographic (XIC) area was used to calculate the relative amount, by dividing to the summation of all XIC from all different glycans and unoccupied with an identical peptide sequence.

#### Phylogenetic tree assembly

We first collected an initial envelope protein sequence dataset comprising 695 yellow fever strains using VIPR (Pickett et al., 2012). We filtered out 335 viral protein sequences with unidentified residues and non-canonical starting and end positions, using as a reference the envelope protein corresponding to the 17D vaccine strain. Next, we filtered out duplicates mapping to the same GenBank identifier, envelope proteins without information regarding their country of origin and envelope proteins corresponding to vaccine strains (except 17D and 17DD vaccines strains). The remaining 281 envelope proteins were further filtered based on pairwise sequence identity (100% sequence identity) and geographical location, keeping only one envelope protein representing identical envelope proteins from different viral strains isolated in the same country. Sequence filtering was performed with CD-HIT (Li and Godzik, 2006). The final non-redundant set of envelope proteins involved 82 yellow fever strains (including vaccine strains 17D and 17DD). Multiple sequence alignment and phylogenetic tree reconstruction was performed using MUSCLE (Edgar, 2004) and PhyML (Guindon et al., 2009) through phylogenetic.fr (Dereeper et al., 2008). Tree visualization was performed with iTol (Letunic and Bork, 2016).

#### Modeling of the glycosylated loop in the envelope protein structure of 17D YFV

We used the modeller plugin (Sali and Blundell, 1993) within Chimera (Pettersen et al., 2004) to model the glycan containing loop in the prefusion state of the 17D envelope dimer structure (missing residues 269 to 272) (Lu et al., 2019). For the glycan, we structurally superimposed the glycosylated N153 found in the envelope protein of Flavivirus Dengue 2 (Rouvinski et al., 2015) to Asn 269 of the 17D YFV envelope protein structure. We chose a loop conformation compatible with the glycosylation that does not lead to steric clashes between the glycan moiety and the envelope dimer.

#### Structural alignment of Flavivirus E proteins complexed with mAbs and the 17D YFV with the modeled glycan

We structurally aligned the experimentally determined monomeric structure of E proteins (complexed with mAbs) from different flaviviruses and the monomeric structure of the 17D E protein (with the modeled glycan) using Chimera (Pettersen et al., 2004). Steric clashes between the modeled glycan and the mAbs (bound to the structurally aligned flavivirus E proteins) were identified using Chimera.

#### B-cell epitope annotation in the envelope protein of YFV

All experimentally known B-cell epitopes on the envelope of YFV strains were extracted from IEDB (Vita et al., 2019). In addition, we predicted B-cell epitopes on the 17D envelope protein using BepiPred-2 (Jespersen et al., 2017).

#### Quantification and statistical analysis

The n number associated with each dataset in the figures indicates the number of biologically independent samples. The number of independent experiments and the measures of central tendency and dispersion used in each case are indicated in the figure legends. Dose-response neutralization curves were fit to a logistic equation by nonlinear regression analysis. Statistical comparisons are indicated in the Figure Legends. All curve-fitting and statistical testing was performed in GraphPad Prism 8.

---

**Table S1.**
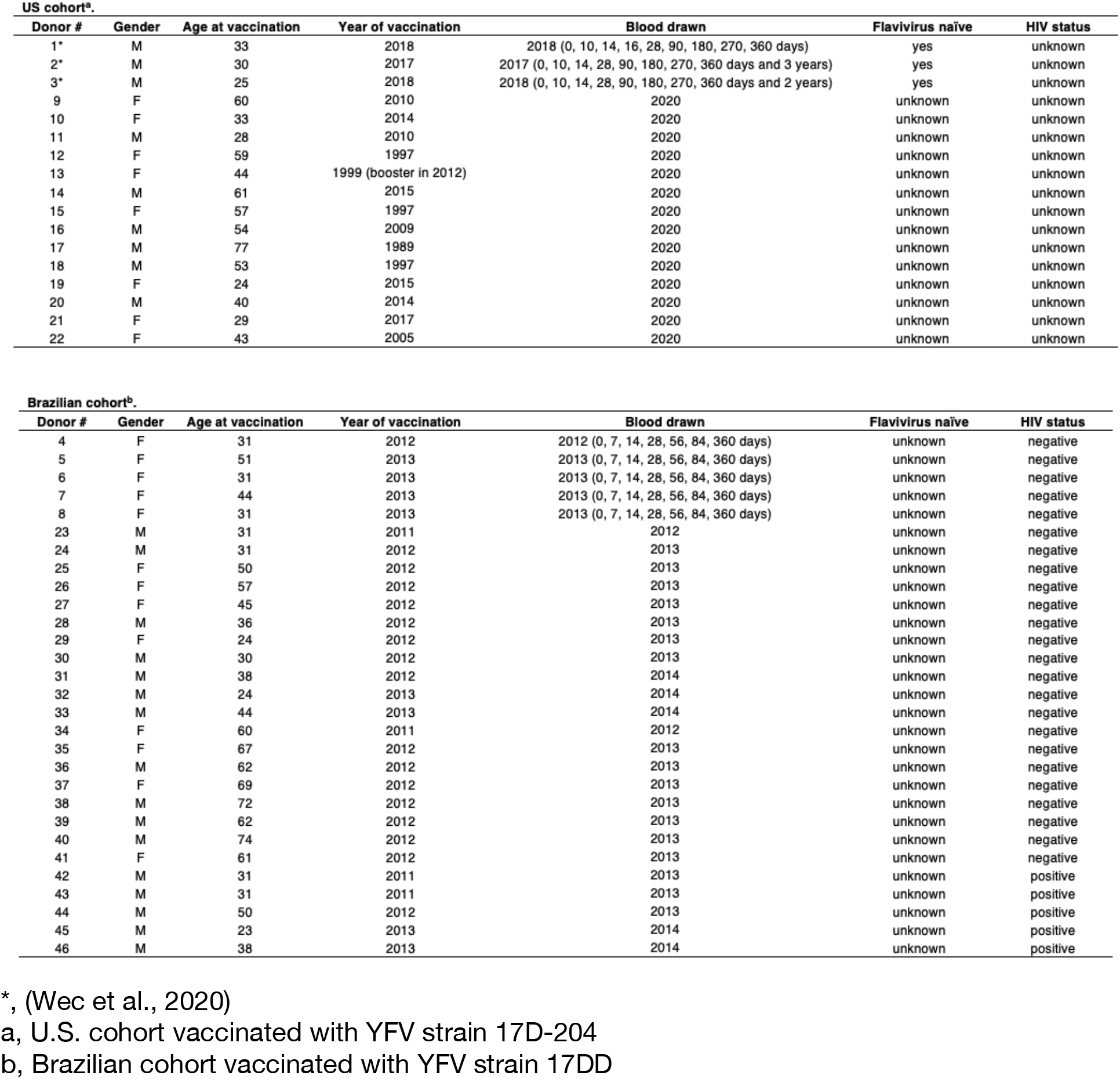
Donor Information.

**Figure S1.**
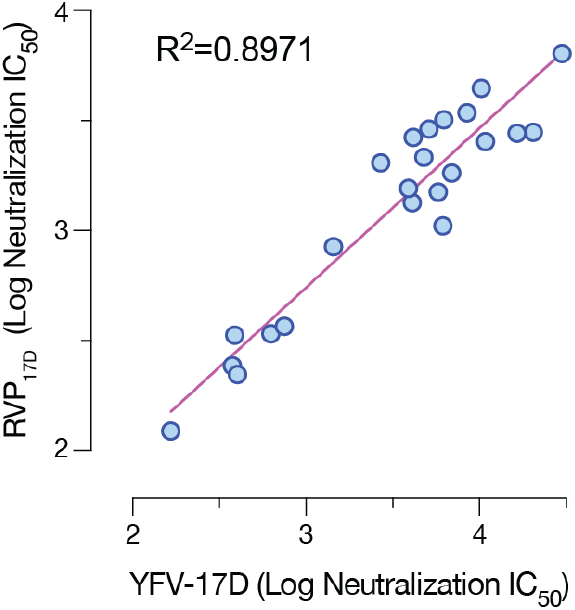
Correlation between serum neutralization titers measured with RVP_17D_ and authentic YFV-17D. Linear regression analyses of neutralization IC_50_ values for RVP_17D_ and authentic YFV-17D for sera from three U.S. vaccinees (see Figure 1A). Means, n=9–12 from three independent experiments).

**Figure S2.**
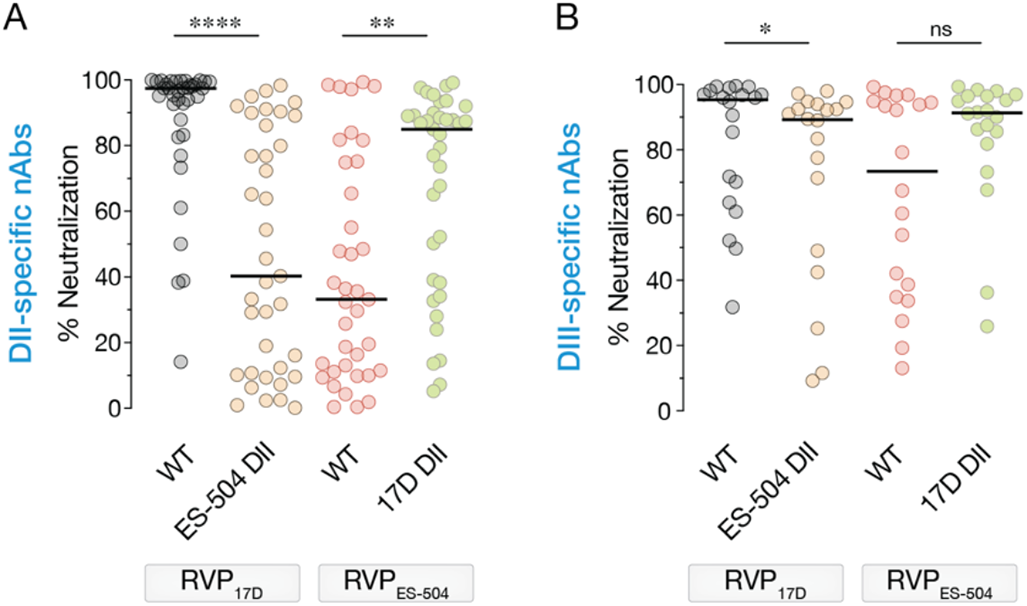
Neutralizing activities of vaccinee nAbs against RVPs bearing 17D/ES-504 domain-swapped E proteins. Neutralizing activities of selected DII- (**A**) and DIII-specific (**B**) nAbs (10 nM) against the indicated RVPs. n=6–9 from 2–3 independent experiments, Parental RVP_17D_ and RVP_ES-504_ controls are from Figure 2. Groups were compared by the Wilcoxon matched-pairs signed-rank test. *, P<0.033. **, P<0.002, ****, P<0.0001. ns, not significant. In all panels, lines indicate group medians.

**Figure S3.**
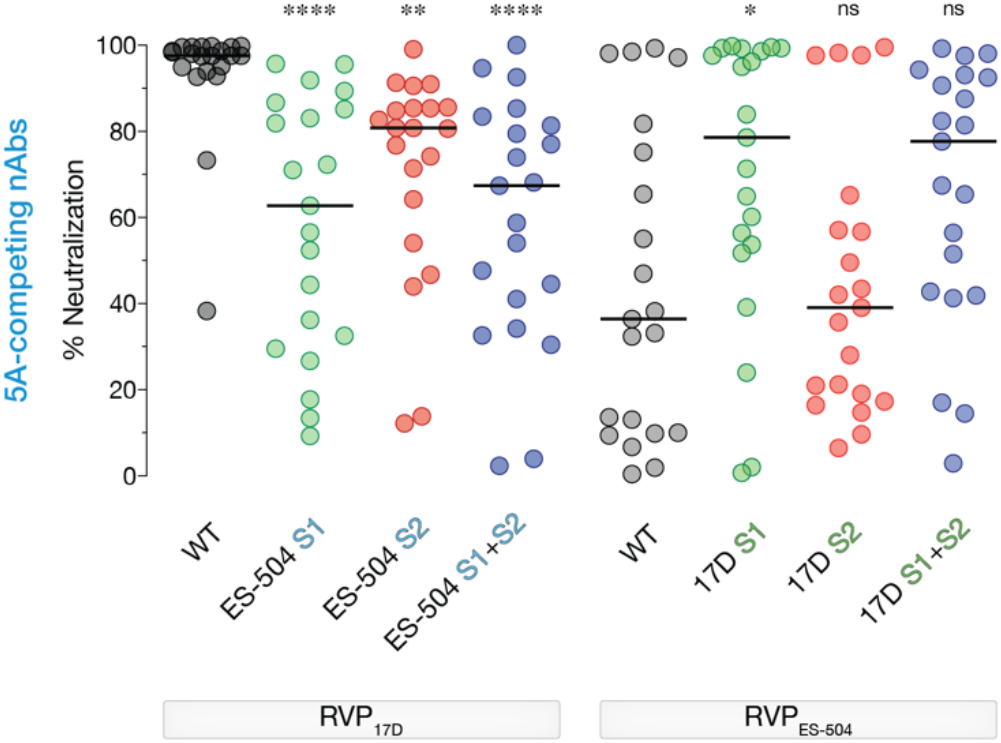
Effects of polymorphisms at Sites 1 and 2 on the neutralizing activities of 5A-competing vaccinee nAbs. Neutralizing activities of selected DII-specific nAbs (10 nM) against the indicated RVPs. Means, n=6–12 from 2–4 independent experiments, Parental RVP_17D_ and RVP_ES-504_ controls are from Figure 2. Groups were compared by two-way ANOVA followed by Tukey’s correction for multiple comparisons. *, P<0.033. ****, P<0.0001. ns, not significant. In all panels, lines indicate group medians.

**Figure S4.**
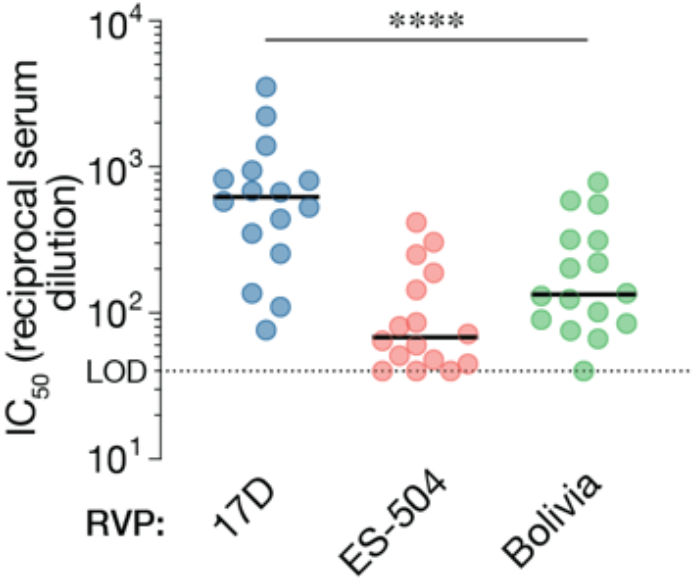
Neutralizing activities of vaccinee sera against RVPs bearing E protein from a genotype 2 South American strain, YFV-Bolivia. Serum neutralizing titers for a U.S. vaccinee cohort against the indicated RVPs. Means, n=6–9 from 2–3 independent experiments. Parental RVP_17D_ RVP_ES-504_ controls are from Figure 1E. Groups were compared by two-way ANOVA followed by Tukey’s correction for multiple comparisons. ****, P<0.0001. In all panels, lines indicate group medians. LOD, limit of detection.

**Figure S5.**
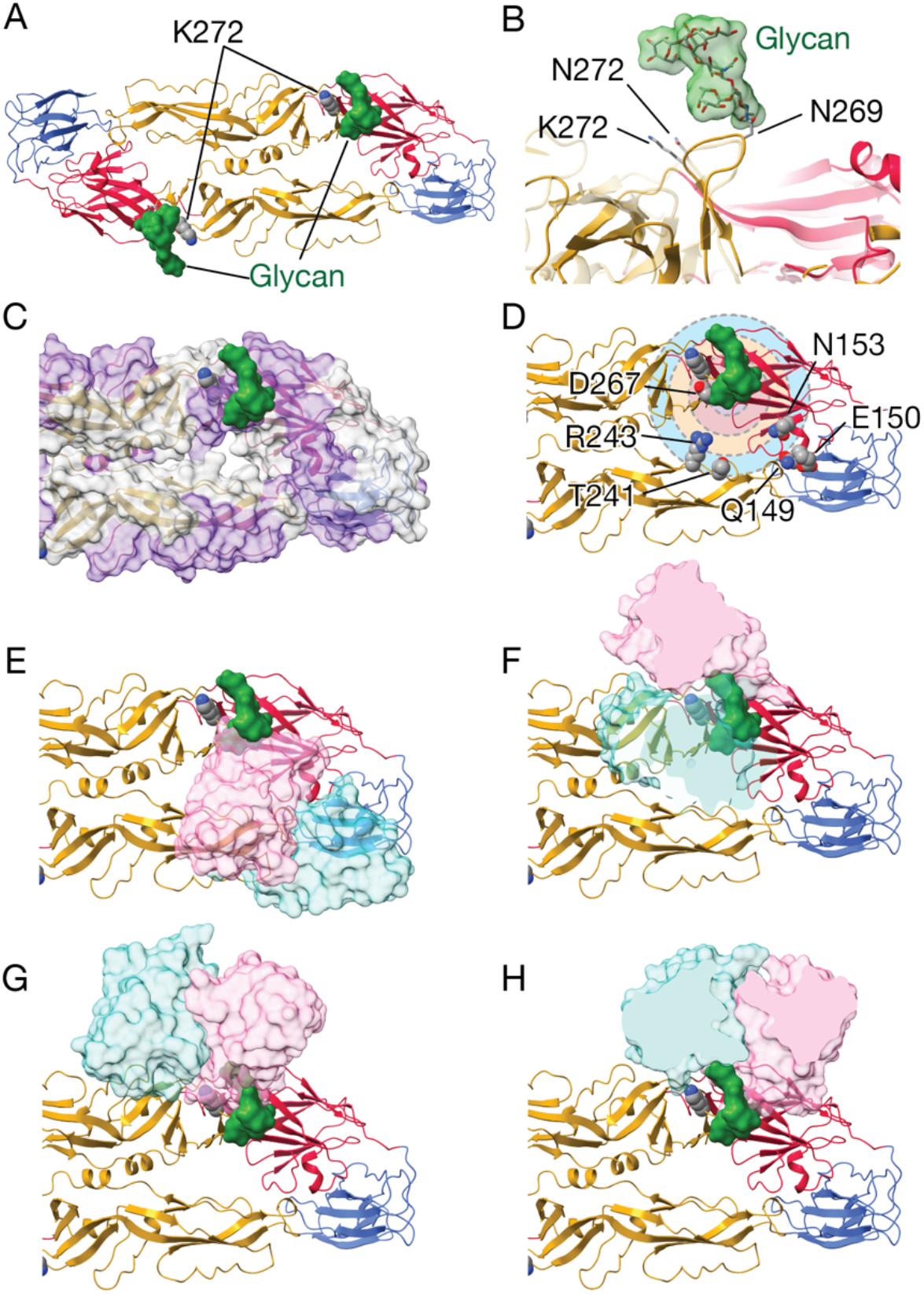
Identification of B-cell epitopes and E:nAb complexes from other flaviviruses in proximity to the *kl* loop in the YFV E protein. (**A**) Dimer structure of YFV-17D E (PDB: 6IW4; Domain I: red, Domain II: yellow, Domain III: blue) with the modeled *kl* loop (residues 267 to 272), including K272 and the glycan moiety (in green) (Lu et al., 2019). The glycan structure corresponds to the experimentally determined glycan moiety observed in the envelope protein of Dengue 2 (glycosylated N153; PDB: 4UTC) (Rouvinski et al., 2015). (**B**) Close-up view of the modeled *kl* loop, the glycan moiety on N269, and K272. Faded structure corresponds to X-ray monomeric structure of 17D E (with Asn at position 272, PDB: 6EPK). (**C–D**) Predicted and known B-cell epitopes in proximity to the *kl* loop. Surface-shaded representation of the YFV-17D E dimer with the modeled *kl* loop (residues 267–272, with K at position 272) and the glycan moiety (in green) is shown. **(C)** Purple colored surface patches correspond to predicted B-cell epitopes. (**D**) Concentric circles (with radius = 5Å, 10Å, 15Å and 20Å) are centered at Cα of glycosylated Asn 269. Residues represented as spheres have been described to be part of B-cell epitopes and contain heavy atoms that fall within the concentric circles: i) D267 (up to 5Å); ii) N153 and R243 (up to 15Å); iii) G102, Q149, E150 and T241 (up to 20Å). (**E–J**) Structural superposition of a monomer of 17D E (ribbon representation) with the modeled *kl* loop and glycan and one E monomer:Fab complex (flavivirus E from each E:Fab complex isnot shown for clarity; Ab Heavy chain: hot pink; Ab Light chain: Cyan; Fab: Grey): (**E**) DENV-2 EDE1 C10 (PDB: 4UT9) (Rouvinski et al., 2015); (**F**) JEV (Japanese Encephalitis virus) 2F2 (PDB: 5YWO) (Qiu et al., 2018); (**G**) ZIKV (Zika virus) SIGN-3C (PDB: 7BU8) (Zhang et al., 2020); (**H**) WNV (West Nile virus) CR4354 (PDB: 3IYW) (Kaufmann et al., 2010). Antibody domains distal to the E protein were clipped out for clarification in panels F–H.

